# A monoclonal antibody that neutralizes SARS-CoV-2 variants, SARS-CoV, and other sarbecoviruses

**DOI:** 10.1101/2021.10.13.464307

**Authors:** Pengfei Wang, Ryan G. Casner, Manoj S. Nair, Jian Yu, Yicheng Guo, Maple Wang, Jasper F.-W. Chan, Gabriele Cerutti, Sho Iketani, Lihong Liu, Zizhang Sheng, Zhiwei Chen, Kwok-Yung Yuen, Peter D. Kwong, Yaoxing Huang, Lawrence Shapiro, David D. Ho

**Affiliations:** Aaron Diamond AIDS Research Center, Columbia University Vagelos College of Physicians and Surgeons, New York, NY 10032, USA; Department of Biochemistry and Molecular Biophysics, Columbia University, New York, NY 10032, USA; State Key Laboratory of Emerging Infectious Diseases, Carol Yu Centre for Infection, Department of Microbiology, Li Ka Shing Faculty of Medicine, The University of Hong Kong, Pokfulam, Hong Kong Special Administrative Region, China; Centre for Virology, Vaccinology and Therapeutics, Hong Kong Science and Technology Park, Hong Kong Special Administrative Region, China; Vaccine Research Center, National Institutes of Health, Bethesda, MD 20892, USA; Department of Microbiology and Immunology, Columbia University Irving Medical Center, New York, NY 10032, USA; Division of Infectious Diseases, Department of Internal Medicine, Columbia University Vagelos College of Physicians and Surgeons, New York, NY 10032, USA

**Author notes:** These authors contributed equally: Pengfei Wang, Ryan G. Casner. School of Life Sciences, Fudan University, Shanghai 200438, China (Pengfei Wang).

## Abstract

The repeated emergence of highly pathogenic human coronaviruses as well as their evolving variants highlight the need to develop potent and broad-spectrum antiviral therapeutics and vaccines. By screening monoclonal antibodies (mAbs) isolated from COVID-19-convalescent patients, we found one mAb, 2-36, with cross-neutralizing activity against SARS-CoV. We solved the cryo-EM structure of 2-36 in complex with SARS-CoV-2 or SARS-CoV spike, revealing a highly conserved epitope in the receptor-binding domain (RBD). Antibody 2-36 neutralized not only all current circulating SARS-CoV-2 variants and SARS-COV, but also a panel of bat and pangolin sarbecoviruses that can use human angiotensin-converting enzyme 2 (ACE2) as a receptor. We selected 2-36-escape viruses *in vitro* and confirmed that K378T in SARS-CoV-2 RBD led to viral resistance. Taken together, 2-36 represents a strategic reserve drug candidate for the prevention and treatment of possible diseases caused by pre-emergent SARS-related coronaviruses. Its epitope defines a promising target for the development of a pan-sarbecovirus vaccine.

## Introduction

Coronaviruses are zoonotic pathogens found in avian and mammalian reservoirs, and seven strains have been found to spillover to humans. Among them, four continually circulate in the human population and only cause mild symptoms of the common cold: 229E and NL63 belong to the *alpha-coronavirus* genus and OC43 and HKU1 belong to the *beta-coronavirus* genus [1]. The other three human coronaviruses are all highly pathogenic and belong to the *beta-coronavirus* genus: severe acute respiratory syndrome coronavirus 2 (SARS-CoV-2), causing the current COVID-19 pandemic, and SARS-CoV, which caused an outbreak 18 years ago, are members of the subgenus *sarbecovirus*; whereas Middle-East respiratory syndrome coronavirus (MERS-CoV) is a member of the *merbecovirus* subgenus [2].

Phylogenetic analysis of the entire genomes grouped SARS-CoV-2 and SARS-CoV with some SARS-related coronaviruses found in bats or pangolins, including bat coronaviruses RaTG13, Rs4231, SHC014, and WIV1, as well as pangolin coronaviruses Pangolin Guangdong and Pangolin Guangxi in the *Sarbecovirus* subgenus [2]. Both SARS-CoV-2 and SARS-CoV express a transmembrane glycoprotein termed spike protein, which mediates viral entry into host cells by engaging ACE2 as the receptor [3,4] and is therefore the primary target of virus-neutralizing antibodies. There is also experimental evidence showing that some of these bat or pangolin viruses could enter into human cells expressing ACE2 [5], indicating their pandemic potential.

SARS-CoV-2 is the causative agent of COVID-19, having infected >238 million people and caused >4.8 million deaths worldwide. Over the past year, several protective vaccines and neutralizing antibody-based therapeutics have become available. However, the emergence of SARS-CoV-2 variants has altered the landscape, threatening the efficacy of these interventions. We and others have shown that some variants such as B.1.351 [6], P.1 [7], B.1.526 [8] and B.1.427/B. 1.429 [9] are more resistant to neutralization by some mAbs, as well as by sera from convalescent patients and vaccinees. As an example, a single mutation, E484K, found in several variants could knock out a class of antibodies binding the receptor binding motif (RBM) on the viral spike [6–8]. Therefore, finding a reagent that can target not only the SARS-CoV-2 mutant variants but also related sarbecoviruses is of utmost importance.

Here we describe the isolation of a mAb that cross-reacts and broadly neutralizes SARS-CoV-2 variants, SARS-CoV, and a panel of bat and pangolin sarbecoviruses. Structural analyses and *in vitro* escape mutation selection indicate that this mAb targeting a highly conserved RBD epitope that could be informative for the development of pan-sarbecovirus vaccines and therapeutics.

## Materials and methods

### Cell lines

HEK293T/17 (cat# CRL-11268) and Vero E6 cells (cat# CRL-1586) were from ATCC, 293T-ACE2 cells were kindly provided by J. Sodroski of Harvard Medical School, and they were cultured in 10% fetal bovine serum (FBS, GIBCO cat# 16140071) supplemented Dulbecco’s Modified Eagle Medium (DMEM, ATCC cat# 30-2002) at 37°C, 5% CO_2_. I1 mouse hybridoma cells (ATCC, cat# CRL-2700) were cultured in Eagle’s Minimum Essential Medium (EMEM, ATCC cat# 30-2003)) with 20% FBS.

### Pseudovirus neutralization assays

Plasmids encoding the single-mutation and the combination of mutations found in SARS-CoV-2 variants were generated by Quikchange II XL site-directed mutagenesis kit (Agilent). Recombinant Indiana vesicular stomatitis virus (VSV) expressing different coronavirus spikes were generated as previously described [10,11]. Briefly, HEK293T cells were grown to 80% confluency before transfection with the spike gene using Lipofectamine 3000 (Invitrogen). Cells were cultured overnight at 37°C with 5% CO_2_, and VSV-G pseudo-typed ΔG-luciferase (G*ΔG-luciferase, Kerafast) was used to infect the cells in DMEM at a multiplicity of infection (MOI) of 3 for 2 hrs before washing the cells with 1X DPBS three times. The next day, the transfection supernatant was harvested and clarified by centrifugation at 300 g for 10 min. Each viral stock was then incubated with 20% I1 hybridoma (anti-VSV-G, ATCC: CRL-2700) supernatant for 1 hr at 37°C to neutralize contaminating VSV-G pseudo-typed ΔG-luciferase virus before measuring titers and making aliquots to be stored at −80°C. Neutralization assays were performed by incubating each pseudovirus with serial dilutions of a mAb and scored by the reduction in luciferase gene expression as previously described [10,11]. Briefly, Vero E6 cells (for SARS-CoV-2 and SARS-CoV) or 293T-ACE2 cells (for bat/pangolin coronaviruses) were seeded in 96-well plates (2 ×10^4^ cells per well). Each pseudovirus was incubated with serial dilutions of a mAb in triplicate for 30 min at 37□°C. The mixture was added to cultured cells and incubated for an additional 16 hrs. Luminescence was measured using Luciferase Assay System (Promega), and IC_50_ was defined as the dilution at which the relative light units were reduced by 50% compared with the virus control wells (virus + cells) after subtraction of the background in the control groups with cells only. The IC_50_ values were calculated using a five-parameter dose-response curve in GraphPad Prism v.8.4.

### Authentic SARS-CoV-2 microplate neutralization

The SARS-CoV-2 viruses USA-WA1/2020 (WA1), hC0V-19/USA/CACDC_5574/2020 (B.1.1.7), hCoV-19/South Africa/KRISP-K005325/2020 (B.1.351), hCoV-19/Japan/TY7-503/2021 (P.1), and hCoV-19/USA/NY-NP-DOH1/2021 (B. 1.526) were obtained from BEI Resources (NIAID, NIH). The viruses were propagated for one passage using Vero E6 cells. Virus infectious titer was determined by an end-point dilution and cytopathic effect (CPE) assay on Vero E6 cells as described previously [10,11]. An end-point dilution microplate neutralization assay was performed to measure the neutralization activity of purified mAbs. Triplicates of each dilution were incubated with SARS-CoV-2 at an MOI of 0.1 in EMEM with 7.5% inactivated FBS for 1 hr at 37°C. Post incubation, the virus-antibody mixture was transferred onto a monolayer of Vero E6 cells grown overnight. The cells were incubated with the mixture for ~70 hrs. CPE was visually scored for each well in a blinded fashion by two independent observers. The results were then converted into percentage neutralization at a given sample dilution or mAb concentration, and the averages ± SEM were plotted using a five-parameter dose-response curve in GraphPad Prism v.8.4.

### SARS-CoV neutralization assay

Antibodies were subjected to successive two-fold dilutions starting from 20 μg/ml. Quadruplicates of each dilution were incubated with SARS-CoV GZ50 strain (GenBank accession no. AY304495) at MOI of 0.01 in DMEM with 2% inactivated FBS for 1 hr at 37°C [12]. After incubation, the virus-antibody mixture was transferred onto a monolayer of Vero E6 cells grown overnight. The cells were incubated with the mixture for 72 hrs. Cytopathogenic effects of viral infection were visually scored for each well in a blinded manner by two independent observers. The results were then converted into the percentage of neutralization at a given monoclonal antibody concentration, and the data were plotted using a five-parameter dose–response curve in GraphPad Prism v.8.4.

### Protein expression and purification

The SARS-CoV-2 and SARS-CoV S2P spike constructs were produced as previously described [3]. The proteins were expressed in HEK293 Freestyle cells (Invitrogen) in suspension culture using serum-free media (Invitrogen) and transfected into HEK293 cells using polyethyleneimine (Polysciences). Cell growths were harvested four days after transfection, and the secreted proteins were purified from supernatant by nickel affinity chromatography using Ni-NTA IMAC Sepharose 6 Fast Flow resin (GE Healthcare) followed by size exclusion chromatography on a Superdex 200 column (GE Healthcare) in 10 mM Tris, 150 mM NaCl, pH 7.4. Spike purity was assessed by SDS-PAGE. 2-36 was expressed and purified as previously described [10]. Fab fragments were produced by digestion of IgGs with immobilized papain at 37 °C for 3 hrs in 50 mM phosphate buffer, 120 mM NaCl, 30 mM cysteine, 1 mM EDTA, pH 7. The resulting Fabs were purified by affinity chromatography on protein A, and purity was assessed by SDS-PAGE.

### ELISA

ELISA detection of mAbs binding to SARS-CoV-2 and SARS-CoV spike trimers was performed as previously described [10]. For the competition ELISA, purified mAbs were biotin-labelled using One-Step Antibody Biotinylation Kit (Miltenyi Biotec) following the manufacturer’s recommendations and purified using 40K MWCO Desalting Column (ThermoFisher Scientific). Serially diluted competitor antibodies (50 μl) were added into spike trimer-precoated ELISA plates, followed by 50 μl of biotinylated antibodies at a concentration that achieves an OD_450_ reading of 1.5 in the absence of competitor antibodies. Plates were incubated at 37□°C for 1 hr, and 100 μl of 500-fold diluted Avidin-HRP (ThermoFisher Scientific) was added into each well and incubated for another 1 hr at 37□°C. The plates were washed with PBST between each of the previous steps. The plates were developed afterwards with 3,3’,5,5’-tetramethylbenzidine (TMB) and absorbance was read at 450 nm after the reaction was stopped. For the ACE2 competition ELISA, 100 ng of ACE2 protein (Abcam) was immobilized on the plates at 4□°C overnight. The unbound ACE2 was washed away by PBST and then the plates were blocked. After washing, 100 ng of S trimer in 50 μl dilution buffer was added into each well, followed by addition of another 50 μl of serially diluted competitor antibodies and then incubation at 37□°C for 1 hr. The ELISA plates were washed four times with PBST and then 100 μl of 2,000-fold diluted anti-strep-HRP (Millipore Sigma) was added into each well for another 1 hr at 37□°C. The plates were then washed and developed with TMB, and absorbance was read at 450 nm after the reaction was stopped.

### Surface plasmon resonance (SPR)

The antibody binding affinity to SARS-CoV-2 and SARS-CoV spike trimers and RBDs was detected by Biacore T200 SPR system (Cytiva). All experiments were performed at 25°C in HBS-EP+ buffer (10 mM HEPES, pH 7.4; 150 mM NaCl; 3.4 mM EDTA; 0.005% (v/v) surfactant P20). The anti-His tag antibodies, diluted at 50 μg/mL in 10 mM sodium acetate, pH 4.5, were immobilized on both the active and reference flow cells surface of the activated CM5 sensor chip using amine coupling method. Approximately 200 RU of His-tagged SARS-CoV-2 and SARS-CoV spike trimers and RBDs were captured onto the chip for the active surface, and anti-His antibody alone served as the reference surface. The antibodies were injected through both flow cells at different concentrations (ranging from 300-1.2 nM in 1:3 successive dilutions) at a flow rate of 30 μL/min for 120 s, followed by a 15 s dissociation step. After each assay cycle, the sensor surface was regenerated with a 30 s injection of 10 mM glycine, pH 1.5, at a flow rate of 30 μL/min. Background binding to reference flow cells was subtracted and antibody binding levels were calculated using Biacore T200 evaluation software (GE Healthcare).

### Cryo-EM grid preparation

Samples for cryo-EM grid preparation were produced by mixing purified spike protein to a final trimer concentration of 0.33 mg/mL with 2-36 Fab in a 1:9 molar ratio, followed by incubation on ice for 1 hr. The final buffer for the 2-36 complex was 10 mM sodium acetate, 150 mM NaCl, pH 5.5. n-Dodecyl β-D-maltoside (DDM) at a final concentration of 0.005% (w/v) was added to the mixtures to prevent preferred orientation and aggregation during vitrification. Cryo-EM grids were prepared by applying 3 μL of sample to a freshly glow-discharged carbon-coated copper grid (CF 1.2/1.3 300 mesh); the sample was vitrified in liquid ethane using a Vitrobot Mark IV with a wait time of 30 s, a blot time of 3 s, and a blot force of 0.

### Cryo-EM data collection and analysis

Cryo-EM data for single particle analysis was collected on a Titan Krios electron microscope operating at 300 kV, equipped with an energy filter and a Gatan K3-BioQuantum direct detection detector, using the Leginon [13] software package. Exposures were taken with a total electron fluence of 41.92 e-/Å2 fractionated over 60 frames, with a total exposure time of 3 seconds. A defocus range of −0.8 to −2.5 μm was used with a magnification of 81,000x, and a pixel size of 1.07 Å.

Data processing was performed using cryoSPARC v2.15 [14]. Raw movies were aligned and dose-weighted using patch motion correction, and the CTF was estimated using patch CTF estimation. Micrographs were picked using blob picker, and a particle set was selected using 2D and 3D classification. The resulting particle set was refined to high resolution using a combination of heterogenous and homogenous refinement, followed by nonuniform refinement. The interface between RBD and 2-36 Fab was locally refined by using a mask that included RBD and the variable domains of the Fab. The final global and local maps were deposited to the EMDB with ID: EMD-24190.

### Model building and refinement

The 2-36-RBD complex model was built starting from template PDB structures 6BE2 (Fab) and 7BZ5 (RBD) using Phenix Sculptor. SARS-CoV-2 S2P spike density was modeled starting with PDB entry 6VXX [15]. Automated and manual model building were iteratively performed using real space refinement in Phenix [16] and Coot [17]. 2-36 Fab residues were numbered according to Kabat numbering scheme. Geometry validation and structure quality assessment were performed using Molprobity [18]. PDBePISA was used to calculate buried surface area [19]. A summary of the cryo-EM data collection, processing, and model refinement statistics is shown in Table S1. The final model was deposited in the PDB with ID 7N5H.

### Structure Conservation Analysis

The conservation of each RBD residue was calculated using the entropy function of the R package bio3d (H.norm column). The calculation was based on the sequence alignment of SARS-CoV, SARS-CoV-2 and SARS-related bat coronavirus. The visualization of sequence entropy was displayed by PyMol version 2.3.2.

### In vitro selection for resistant mutations against mAb 2-36

SARS-Cov-2 isolate USA-WA1/2020 was mixed with serial five-fold dilutions of 2-36 antibody at MOI 0.2 and incubated for 1 hr. Following incubation, the mix was overlaid on 24-well plate to a final volume of 1mL. the plates were incubated at 37°C for 70 hrs till CPE was complete (100%) in virus control wells bearing no antibody. At this time, all wells were scored to determine the 50% inhibition titer (EC50) and supernatant collected from this well was used for subsequent round of selection. Passaging continued till the virus was able to form CPE in the presence of 50 μg/mL of 2-36 antibody. At this point, the resulting supernatant was collected, and RNA was extracted using QiaAMP Viral RNA kit (Qiagen). cDNA was obtained using Superscript IV enzyme (Thermo Scientific). Amplification of spike gene from cDNA was performed using nested PCR and sequenced using Sanger sequencing (Genewiz). Multiple clones from limiting dilution nested PCR were sequenced to confirm the dominant mutants in the pool of the resulting progeny viruses and a percentage of their prevalence was calculated from total number sequenced. For passage 4, 9 and 12, a total of 20, 10 and 10 clones were sequenced respectively to confirm the mutations.

### Data availability

The cryo-EM structure of antibody 2-36 in complex with prefusion SARS-CoV-2 spike glycoprotein has been deposited in the PDB ID: 7N5H and EMDB ID: 24190.

## Results

### Identification of a SARS-CoV-2 and SARS-CoV cross-reactive mAb from a COVID-19 patient

By single B-cell sorting and 10X Genomics sequencing, we have previously recovered 252 mAb sequences from five COVID-19 patients and isolated 19 potent neutralizing mAbs [10] targeting different epitopes on SARS-CoV-2 spike [10,20–22]. To identify mAbs with broad reactivity against other coronaviruses, we screened these 252 mAb transfection supernatants for neutralization against SARS-CoV pseudovirus in the same way we did for SARS-CoV-2 pseudovirus. Although about one fifth of mAbs showed neutralization activities against SARS-CoV-2 [10], only one, 2-36, had an appreciable neutralization potency against SARS-CoV (Figure 1A).

**Figure 1.**
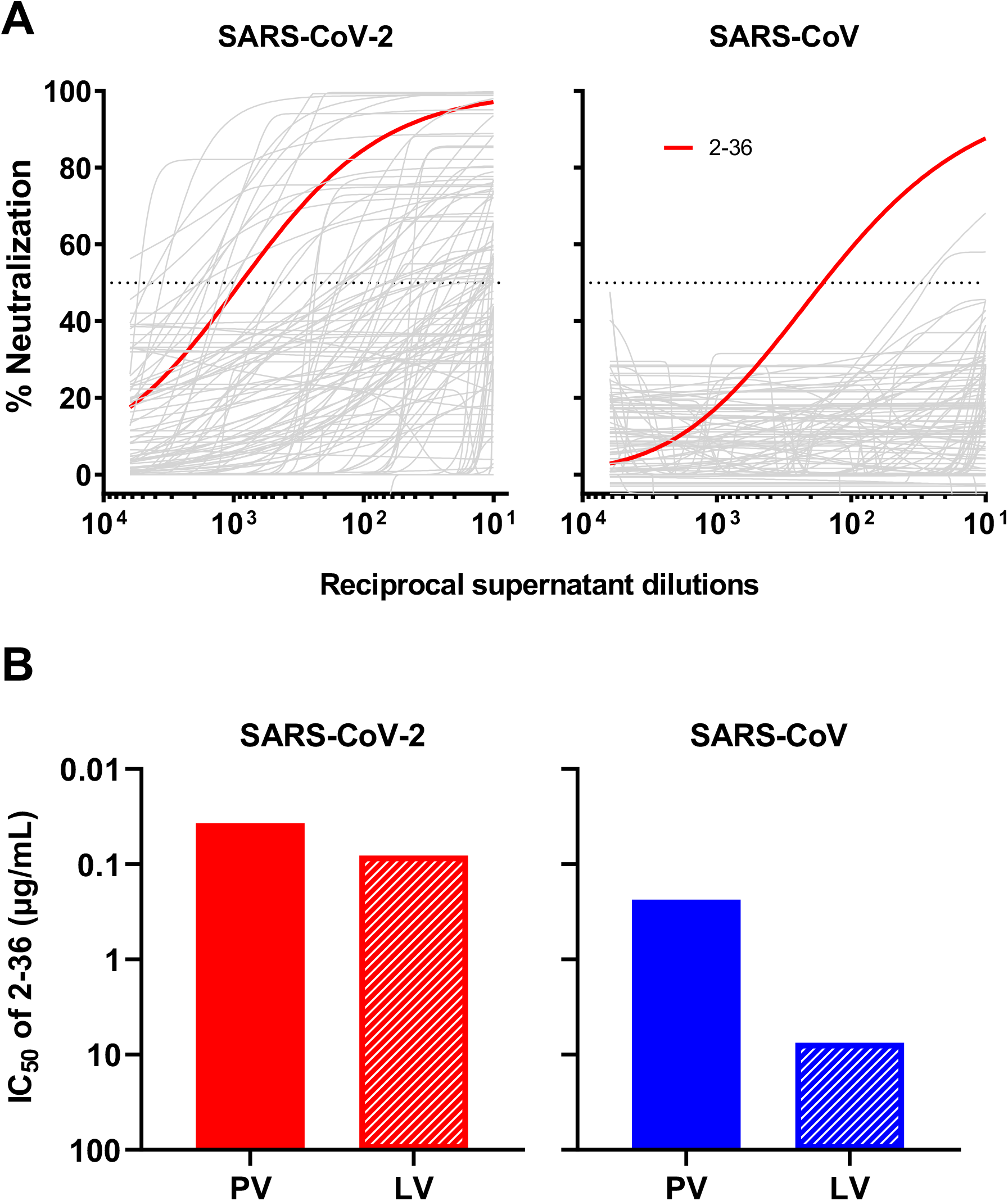
2-36 neutralizes both SARS-CoV-2 and SARS-CoV. **(A)** Screening of mAb transfection supernatant for neutralizing activity against SARS-CoV-2 and SARS-CoV pseudoviruses. **(B)** 2-36 neutralization IC_50_ (μg/mL) against SARS-CoV-2 and SARS-CoV pseudoviruses (PV) and live viruses (LV).

We then focused on 2-36, by carefully characterizing it on SARS-CoV-2 and SARS-CoV with purified antibody. It neutralized SARS-CoV-2 pseudovirus at IC_50_ ~0.04 μg/mL and the authentic virus (WA1 strain) at IC_50_ ~0.1 μg/mL. Its potency versus SARS-CoV was lower, with IC_50_ ~0.2 μg/mL against the pseudovirus and IC_50_ ~7.5 μg/mL against the authentic virus (GZ50 strain) (Figure 1B). While 2-36 could bind to both SARS-CoV-2 and SARS-CoV spike trimer (Supplementary Figure 1), its binding affinity to SARS-CoV-2 spike measured by SPR was much higher than that to SARS-CoV (Supplementary Figure 2A). As our previous study already showed 2-36 is an RBD-directed mAb [10], we also compared its binding affinity to SARS-CoV-2 and SARS-CoV RBDs, and observed similar trend as seen for the spike proteins (Supplementary Figure 2B), with higher affinity found for SARS-CoV-2 RBD.

### Cryo-EM structure of 2-36 in complex with SARS-CoV-2 and SARS-CoV spikes

To gain insight into the epitope of 2-36, we first evaluated its competition with other mAbs in binding to SARS-CoV-2 spike trimer by ELISA. Two mAbs isolated from SARS patients and showed cross-reactivity against SARS-CoV-2 by targeting distinct epitopes, CR3022 [23] and S309 [24], together with our SARS-CoV-2 mAb targeting the RBM, 2-4 [10], were used in the competition experiments. 2-36 binding to SARS-CoV-2 spike was inhibited by CR3022, but not by S309 or 2-4 (Supplementary Figure 3A), suggesting 2-36 targeted a region similar to the “highly conserved cryptic epitope” of CR3022 [23]. To further investigate the molecular nature of the 2-36 epitope, we determined cryo-EM structures for 2-36 Fab in complex with both SARS-CoV-2 and SARS-CoV spike (S2P-prefusion-stabilized trimers). In the SARS-CoV-2 structure, a single predominant population was observed wherein three 2-36 Fabs were bound per spike in a 3-RBD-up conformation (Figure 2A and Supplementary Figure 4). In the SARS-CoV structure, two 3D classes were observed: one Fab bound per spike in a 1-RBD-up conformation and two Fabs bound per spike in a 2-RBD-up conformation (Figure 2B and Supplementary Figure 5). For the SARS-CoV-2 complex structure, local refinement provided side chain resolution for much of the interface, allowing construction of a molecular model.

**Figure 2.**
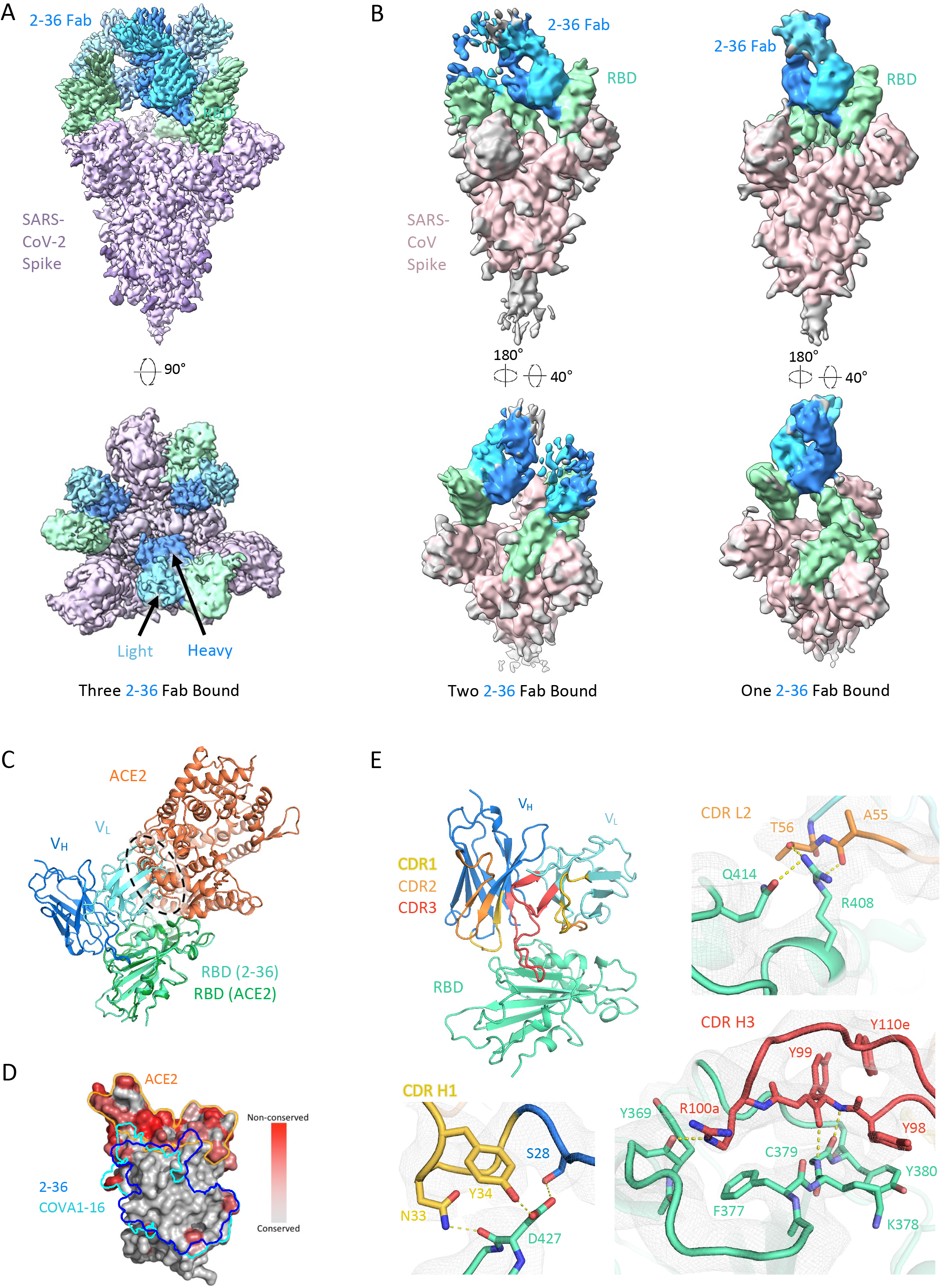
Cryo-EM structure of 2-36 in complex with SARS-CoV-2 and SARS-CoV Spike. **(A)** Cryo-EM reconstruction of 2-36 Fab in complex with the SARS-CoV-2 S trimer at 3.4 Å. RBD is colored in green, the 2-36 Fab heavy chain in darker blue, the light chain in lighter blue, and the rest of the spike is colored in light purple with glycans in darker purple. **(B)** Cryo-EM reconstructions of 2-36 Fab in complex with the SARS-CoV S trimer reveal two major classes: one 2-36 Fab bound to spike with 1-RBD up, and two 2-36 Fabs bound to spike with 2-RBD up. Reconstructions are shown in two different orientations. **(C)** The 2-36 interface model superposed onto an ACE2-RBD complex model (pdb: 6M0J) shows an ACE2 clash with the light chain. **(D)** Conservation analysis on the RBD among SARS-CoV-2, SARS-CoV-1, and bat and pangolin sarbecoviruses show 2-36 binding site is highly conserved, with regions of high conservation in grey and low conservation in red. The 2-36 epitope is outlined in dark blue, and the epitope for COVA1-16 is outlined in cyan. **(E)** The interface model depicted in ribbon representation, with the CDR1 loops in gold, CDR2 loops in orange, and CDR3 loops in red. The interface residue contacts are show for CDR H1, H3, and L2 complemented with the electron density map for a 4.1 Å local reconstruction.

Antibody 2-36 recognizes a region on the ‘inner-side’ of RBD that is buried in the RBD-down conformation of the spike [6]. Thus, 2-36 recognizes RBD only in the up position. This is similar to the antibody epitopes previously defined for antibodies CR3022 [23] and COVA1-16 [25]. The positioning of the 2-36 light chain causes a clash with ACE2 in the spike-ACE2 complex (Figure 2C), which is consistent with competition ELISA data (Supplementary Figure 3B), suggesting that blockage of receptor binding likely accounts for neutralization by 2-36. The 2-36 epitope on RBD is highly conserved among sarbecoviruses that utilize ACE2 for binding, including SARS-CoV-2, SARS-CoV, and a panel of SARS-related coronaviruses (Figure 2D). For example, 24 out of 27 amino acids that contact 2-36 are identical based on the sequence alignment of SARS-CoV-2 Wuhan-Hu-1 and SARS-CoV BJ01 (Supplementary Figure 6), consistent with a similar binding mode between SARS-CoV-2 and SARS-CoV (Figure 2A and 2B).

The interaction of 2-36 with RBD is dominated by CDR H3 (472 Å^2^ buried in the interface), with minor contributions from CDR H1 (176 Å^2^ buried) and CDR L2 (145 Å^2^ buried) (Figure 2E). CDR H3 forms most of the interactions by interacting with a loop on RBD comprising residues 369-385. Residue 99 in the CDR H3 forms backbone hydrogen bonds with RBD residue 379 to extend an RBD β-sheet, much like antibodies C118 and C022 [26] (Figure 2E, bottom right). For 2-36, interactions were primarily hydrophobic, with CDR H3 residues Tyr98, Tyr99, the aliphatic chain of Arg100a, and Tyr100e all making significant contacts with hydrophobic residues on RBD.

### 2-36 broadly neutralizes SARS-CoV-2 variants and SARS-related sarbecoviruses

Given the high conservation of the 2-36 epitope, we went on to test its breadth on the recent emerging SARS-CoV-2 variants first. Our previous studies already showed 2-36 retained its activities against SARS-CoV-2 variants B.1.1.7, B. 1.351 [6], P.1 [7] and B.1.526 [8], on both authentic viruses and pseudoviruses (replotted in Figure 3A). Here we assessed 2-36 activity on more variants, including pseudoviruses representing the combination of key spike mutations of B.1.427/B.1.429, R.1, B. 1.1.1, B. 1.525, B. 1.617.1, B.1.617.2 and B.1.1.7 with E484K, as well as many pseudoviruses with single spike mutations which are naturally circulating in COVID-19 patients with high frequency and located in the N-terminal domain, RBD, or S2. As shown in Figure 3A, 2-36 could neutralize all the variants tested and maintained its potency against most of the variants with IC_50_ below 0.1 μg/mL.

**Figure 3.**
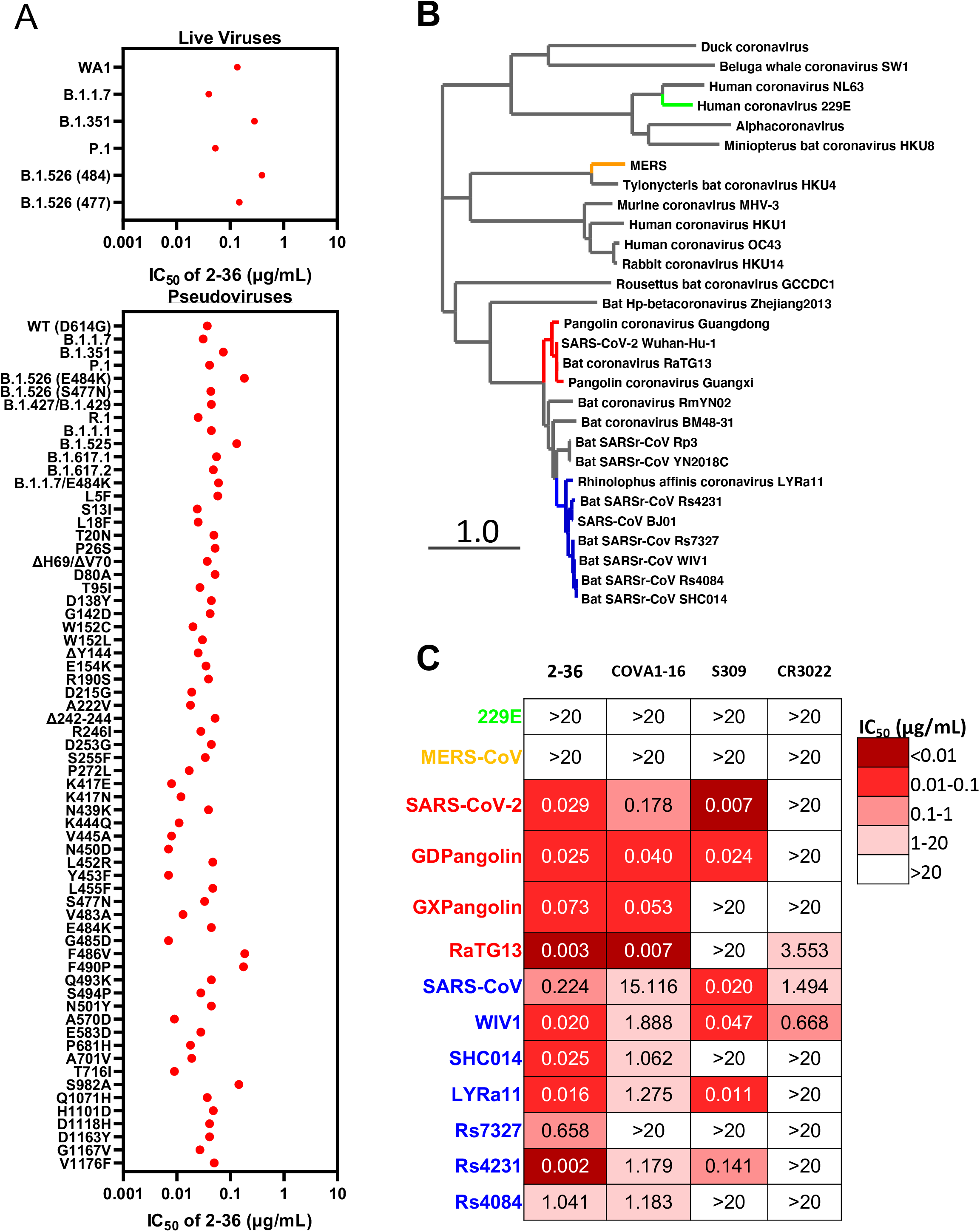
2-36 neutralizes SARS-CoV-2 variants and SARS-CoV-like coronaviruses. **(A)** 2-36 neutralization IC_50_ (μg/mL) against SARS-CoV-2 variants. **(B)** Phylogenetic tree of SARS-CoV-2- and SARS-CoV-related lineages and other coronaviruses constructed via MEGA7 and maximum likelihood analysis of spike amino acid sequences extracted from the NCBI and GISAID database. Representative viruses selected for further testing are denoted in color same as in **(C)**. **(C)** Heatmap showing the neutralization IC_50_ values of the indicated antibodies against SARS-CoV-2, SARS-CoV and their related sarbecoviruses.

We further explored 2-36’s potential as a broadly neutralizing antibody against bat or pangolin coronaviruses (Figure 3B) in the SARS-CoV-2-related lineage (bat coronavirus RaTG13, pangolin coronavirus Guangdong and pangolin coronavirus Guangxi) and SARS-CoV-related lineage (bat WIV1, SHC014, LYRa11, Rs7327, Rs4231 and Rs4084), each of which can use human ACE2 as receptor [2,5]. For comparison, we also tested COVA1-16, CR3022 and S309 in parallel with 2-36. As indicated by the heatmap in Figure 3C and neutralization curves in Supplementary Figure 7, 2-36 could neutralize all these sarbecoviruses. COVA1-16, with a similar epitope as 2-36 (Figure 2D), neutralized most of the viruses but with lower potency. S309 and CR3022 could only neutralize part of this panel of viruses. None of the four antibodies could neutralize MERS (*merbecovirus*) or 229E (*alpha-coronavirus*), reflecting their longer genetic distance from SARS-CoV-2 and SARS-CoV (Figure 3B and C).

### *In vitro* selection of 2-36 escape virus

To test whether SARS-CoV-2 can escape from 2-36 neutralization, we co-incubated the authentic SARS-CoV-2 (WA1 strain) with serially diluted 2-36, and repeatedly passaged the virus from wells showing 50% cytopathic effect (CPE) again in serial dilutions of the antibody. 2-36 retained it neutralization activities on the serially passaged viruses until passage 10 (Supplementary Figure 8), and then at passage 12, the virus became resistant to the antibody (Figure 4A). Sequence analyses of the passage 12 virus revealed four single point spike mutations (T284I, K378T, H655Y, V1128A), all of which were found at 100% frequency. And when we went back to sequence the viruses from earlier passages, only H655Y, which has also been found in other circulating variants such as P.1 [27,28], and V1128A were found. The T284I and K378T mutations appeared only in the 2-36-resistant virus (Figure 4B). We localized the selected mutations in the model of 2-36 in complex with SARS-CoV-2 S trimer (Figure 4C), only K378 resided in RBD and showed a strong van der Waals contact with Y98 of the 2-36 heavy chain, while all the other three residues were distal from the antibody interface. Indeed, when these mutations were introduced into pseudoviruses and tested for their sensitivity to 2-36, only K378T alone or in combination with the other mutations was found to be resistant to 2-36, whereas viruses with T284I, H655Y, or V1128A alone remained sensitive (Figure 4D). Interestingly, although the K378 position in SARS-CoV-2 spike can be mutated to other residues at very low frequency, we could not find any K378T mutation circulating in patient viruses to date (Figure 4E), further demonstrating the conserved nature of the region recognized by 2-36.

**Figure 4.**
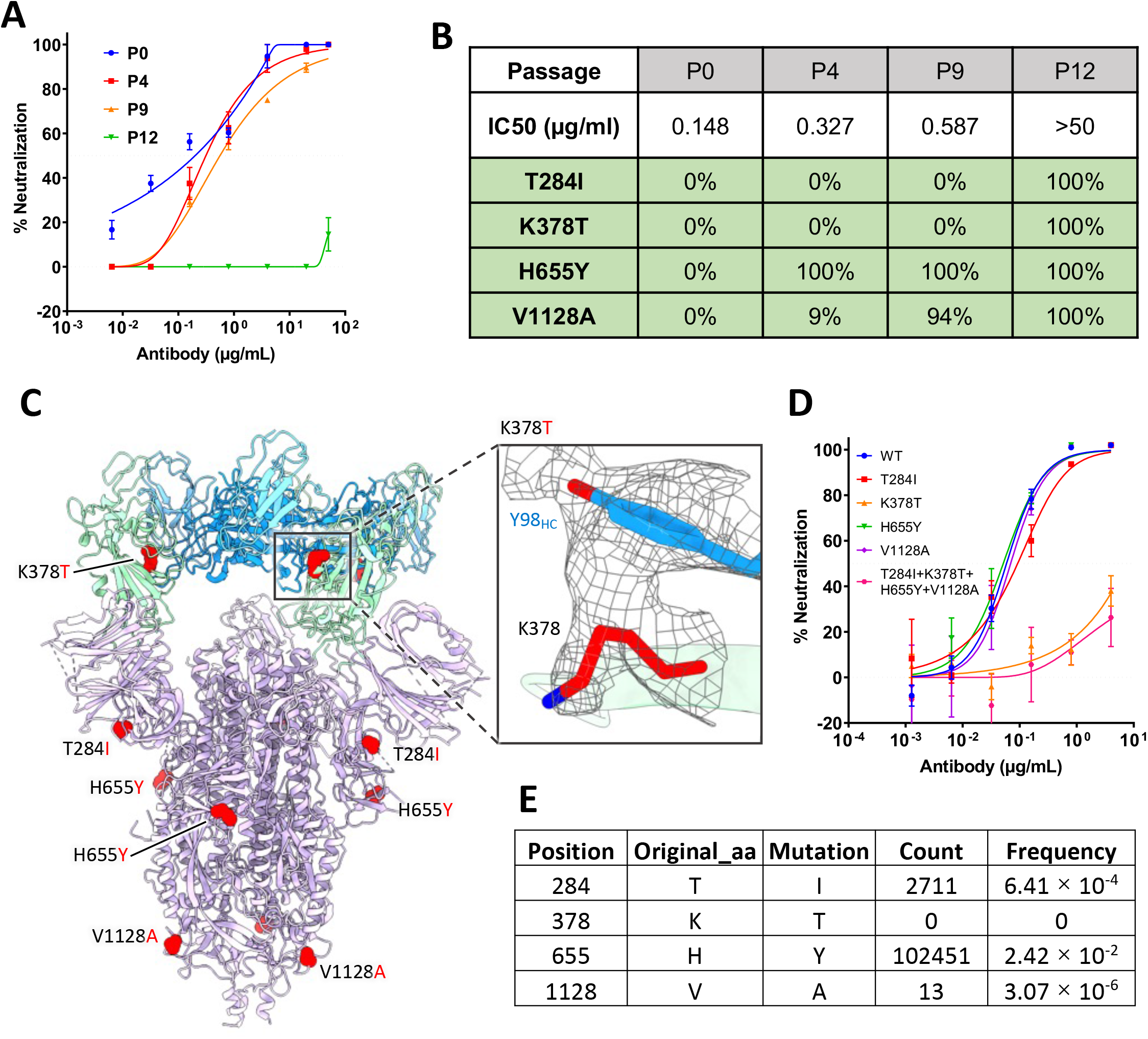
In vitro selection of 2-36 escape viruses. **(A)** Neutralizing activity of 2-36 against viruses at different passages. **(B)** Spike mutations found in viruses at different passages **(C)** Model of 2-36 in complex with the SARS-CoV-2 S trimer highlighting mutations in red. CDR H3 Tyrosine 98 makes van der Waals contacts with RBD residue 378 as shown as electron density map mesh in the subpanel. **(D)** The selected mutations were introduced into pseudoviruses and then tested for neutralization sensitivity to 2-36. **(E)** The frequency of the selected mutations in circulating in infected patients (data updated to Oct 13^th^, 2021).

## Discussion

We have seen the repeated emergence of novel highly pathogenic human coronaviruses that seriously threaten public health and the global economy in the last two decades. While we are still in the midst of the COVID-19 pandemic, other animal coronaviruses with the potential to cross the species-barrier to infect humans need to be considered as potential threats in the future. Therefore, the development of potent and broad-spectrum antiviral interventions against current and emerging coronaviruses is critical. Here, in this study, we described a mAb, 2-36, that exhibited broadly neutralizing activity against not only SARS-CoV-2 variants and SARS-CoV but also a panel of related sarbecoviruses.

Several other studies have also reported the discovery of broadly neutralizing mAbs against sarbecoviruses in addition to the controls used in this study, COVA1-16 and S309. For example, H014 [29] and Ey6a [30] were reported to neutralize SARS-CoV and SARS-CoV-2. Wec et al isolated several antibodies from a SARS survivor that neutralized SARS-CoV, SARS-CoV-2, and the bat SARS-CoV-like virus WIV1 with modest potency [31]. In addition, antibodies C118 and C022 were also shown to neutralize another bat SARS-CoV-like virus SHC014 [26]. Compared to these studies, we have included a much larger panel of viruses for testing the neutralization breadth of 2-36, with 4 viruses in the SARS-CoV-2-related lineage and 7 viruses in the SARS-CoV-related lineage (Figure 3C). Very recently, additional antibodies with broad activities, such as S2X259 [32] and DH1047 [33], targeting a similar surface on the inner side of RBD as 2-36 (Supplementary figure 9), have been reported, again highlighting that the conserved region of the inner face of RBD could represent a target for a pan-sarbecovirus vaccine.

Apart from these RBD-directed mAbs, antibodies targeting the conserved S2 stem-helix region of the coronavirus spike fusion machinery have been shown to possess even broader reactivity against more beta-coronaviruses including MERS [34,35]. However, their relatively low neutralization potency may limit their clinical utility. Using an in vitro affinity maturation strategy, CR3022 has been re-engineered to neutralize SARS-CoV-2 more potently [36]. Similarly, Rappazzo et al engineered a SARS-CoV-2 and SARS-CoV mAb into a better version, ADG-2, with enhanced neutralization breadth and potency [37]. Perhaps 2-36 and S2-specific mAbs could be improved in the same way without sacrificing neutralization breadth.

The cryo-EM structures of 2-36 in complex with both SARS-CoV-2 and SARS-CoV spike trimers revealed a highly conserved epitope. This region on the inner side of RBD is also targeted by several other mAbs [25,26,30]. The structural information from these studies of broadly neutralizing antibodies isolated from natural infection could be valuable in guiding immunogen design for the development of pan-sarbecovirus vaccines. Such vaccines and 2-36-like broadly neutralizing antibodies could be developed and stockpiled to prevent or mitigate future outbreaks of sarbecoviruses.

## Acknowledgements

We thank Bob Grassucci and Chi Wang for help with cryo-EM data collection at the Columbia University cryo-EM Center at the Zuckerman Institute. This study was supported by funding from Andrew & Peggy Cherng, Samuel Yin, Barbara Picower and the JPB Foundation, Brii Biosciences, Roger & David Wu, the Bill and Melinda Gates Foundation, and the SAVE program from the NIH. Support was also provided by the Intramural Program of the Vaccine Research Center, National Institute of Allergy and Infectious Diseases, National Institutes of Health, and Health@InnoHK, Innovation and Technology Commission, the Government of the Hong Kong Special Administrative Region.

## Author contributions

The study was conceptualized by D.D.H. The biological experiments and analyses were carried out by P.W., M.S.N., J.Y., M.W., J.F.-W.C., S.I., L.L., Z.C., K-Y.Y., and Y.H. The structural experiment and analysis were carried out by R.G.C., G.C., Y.G., Z.S., P.D.K., and L.S. The manuscript was written by P.W., R.G.C., L.S., and D.D.H. and reviewed, commented, and approved by all the authors.

---

**Supplementary Figure 1.** Binding of 2-36 to SARS-CoV-2 and SARS-CoV spike as determined by ELISA.

**Supplementary Figure 2.** 2-36 binding affinity to SARS-CoV-2 and SARS-CoV (A) spike or (B) RBD as measured by SPR.

**Supplementary Figure 3.** 2-36 binding to SARS-CoV-2 spike is inhibited by CR3022; 2-36 inhibits hACE2 binding to SARS-CoV-2 spike.

**Supplementary Figure 4. Cryo-EM data processing for antibody 2-36 in complex with the SARS-CoV-2 S trimer.**

**(A)** Representative micrograph, power spectrum, and contrast transfer function (CTF) fit.

**(B)** Representative 2D class averages showing spike particles.

**(C)** Global consensus refinement Fourier Shell Correlation (FSC) curve and particle projection viewing angle distribution.

**(D)** Local focused refinement FSC curve and viewing direction distribution.

**(E)** Local resolution estimation mapped on surface density for global refinement.

**(F)** Local resolution estimation mapped on surface density for local refinement.

**Supplementary Figure 5. Cryo-EM data processing for antibody 2-36 in complex with the SARS-CoV S trimer.**

**(A)** Representative micrograph, power spectrum, and contrast transfer function (CTF) fit.

**(B)** Representative 2D class averages showing spike particles.

**(C)** Global consensus refinement Fourier Shell Correlation (FSC) curve and particle projection viewing angle distribution.

**(D)** Local resolution estimation mapped on surface density for global consensus refinement.

**Supplementary Figure 6. Sequence alignment for SARS-CoV-2 and SARS-CoV RBD binding interface of 2-36.** The dots represent the conserved residues in SARS-CoV-1 compared to SARS-CoV-2. The interface residues are colored in red, residues form hydrogen bond with 2-36 are labeled by underline.

**Supplementary Figure 7.** 2-36 Neutralizes SARS-like coronaviruses using hACE2.

**Supplementary Figure 8.** 2-36 neutralization IC_50_ (μg/mL) on the serially passaged virus.

**Supplementary Figure 9.** Structural comparison between antibody 2-36 in complex with SARS CoV-2 RBD and other published antibody structures.

(A) Molecular models for COVA1-16 (dark blue), 2-36 (teal), and S2X259 (orange), aligned based on RBD, all bind to a similar face on the inner part of RBD.

(B) Close up of the aforementioned antibody CDRH3 loops all target the same beta-strand on the surface of the inner face of RBD.

(C) Comparison of binding footprints for published broadly neutralizing antibodies that bind to the same inner face of RBD.

**Supplementary Table 1.** Cryo-EM data collection, processing, and model refinement and validation statistics. Related to Figures 2 and 4.

