## Supplementary figures for "A monoclonal antibody that neutralizes SARS-CoV-2 variants, SARS-CoV, and other sarbecoviruses"

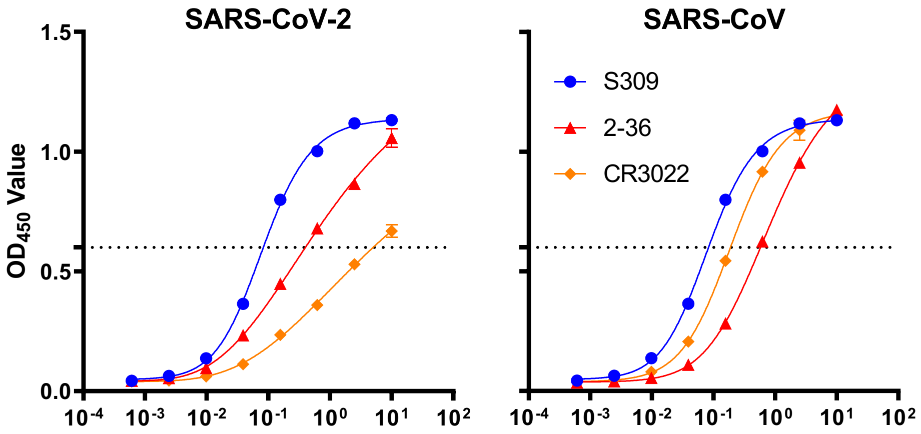


**Supplementary Figure 1.** Binding of 2-36 to SARS-CoV-2 and SARS-CoV spike as determined by ELISA.

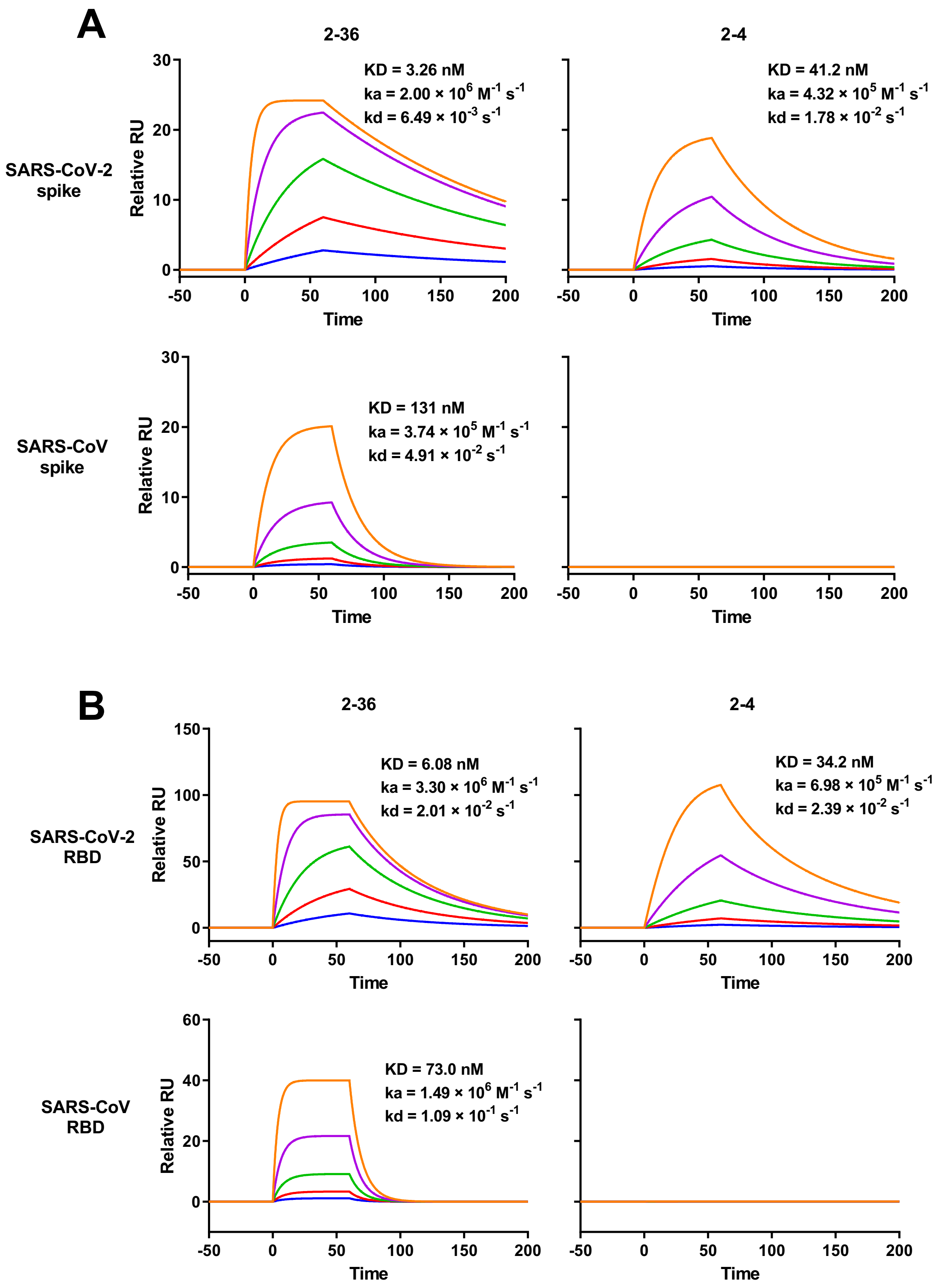


**Supplementary Figure 2.** 2-36 binding affinity to SARS-CoV-2 and SARS-CoV (a) spike or (b) RBD as measured by SPR.

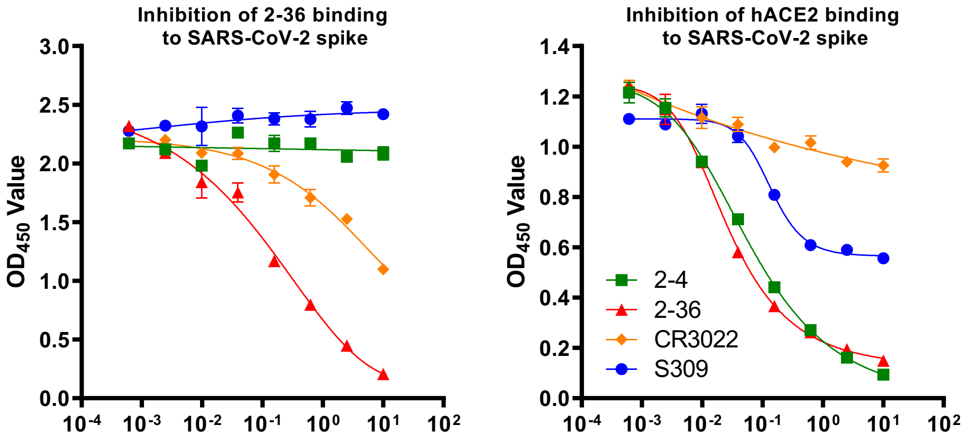


**Supplementary Figure 3.** 2-36 binding to SARS-CoV-2 spike is inhibited by CR3022; 2-36 inhibits hACE2 binding to SARS-CoV-2 spike.

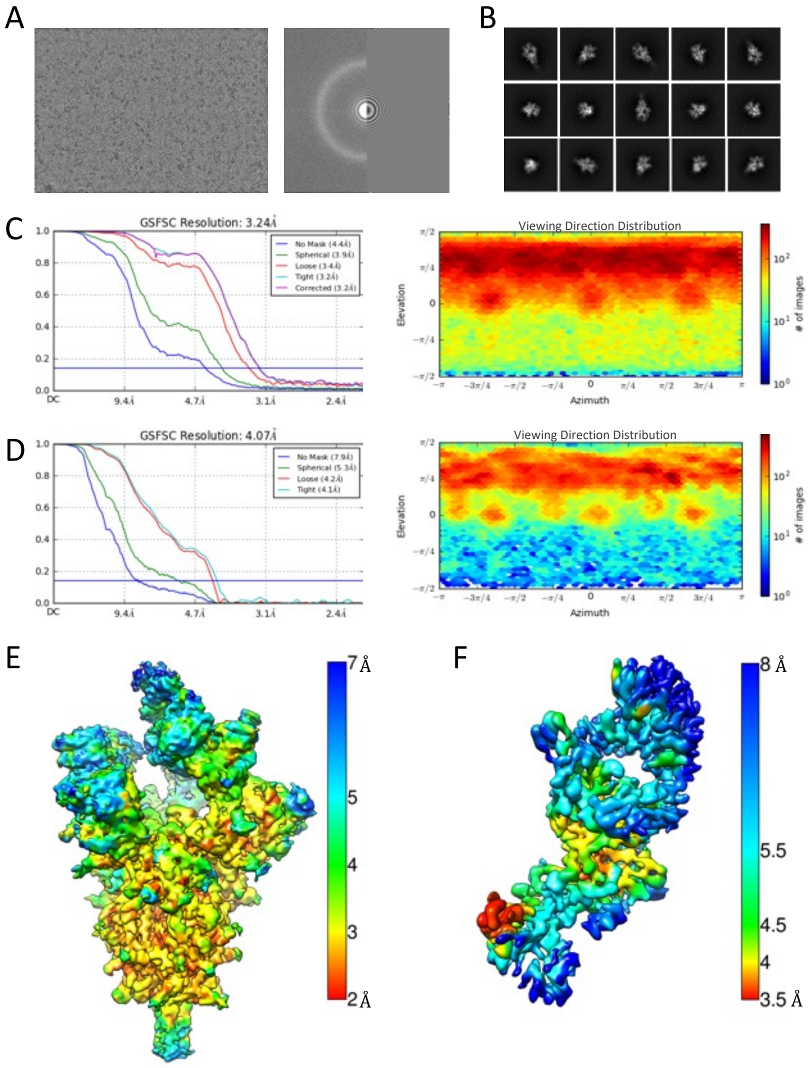


**Supplementary Figure 4. Cryo-EM data processing for antibody 2-36 in complex with the SARS-CoV-2 S trimer.**

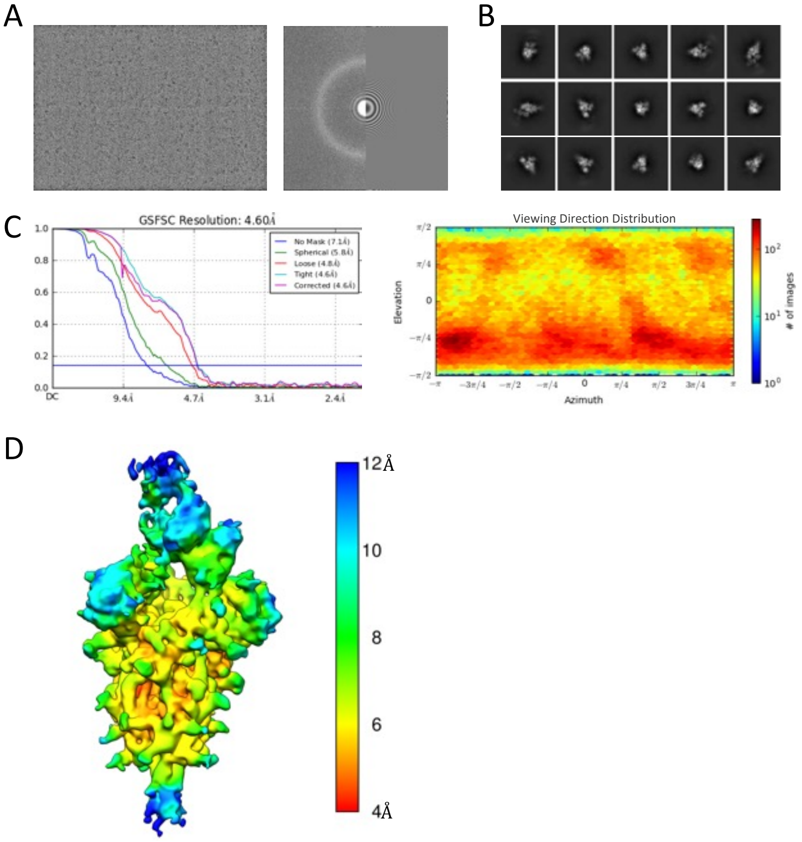


**Supplementary Figure 5. Cryo-EM data processing for antibody 2-36 in complex with the SARS-CoV S trimer.**

```
                           350                      375                      400
SARS-CoV-2 Wuhan-Hu-1  FASVYAWNRKRISNCVADYSVLYNSASFSTFKCYGVSPTKLNDLCFTNVYADSFVIRGDEVRQIA
SARS-CoV BJ01          .P.....E.................TF..........A........S........VK..D.....

                                     425                      450                      475
SARS-CoV-2 Wuhan-Hu-1  PGQTGKIADYNYKLPDDFTGCVIAWNSNNLDSKVGGNYNYLYRLFRKSNLKPFERDISTEIYQAGST
SARS-CoV BJ01          .....V............M...L...TR.I.ATST.....K..YL.HGK.R.......NVPFSPDGK
```

**Supplementary Figure 6. Sequence alignment for SARS-CoV-2 and SARS-CoV RBD binding interface of 2-36.**

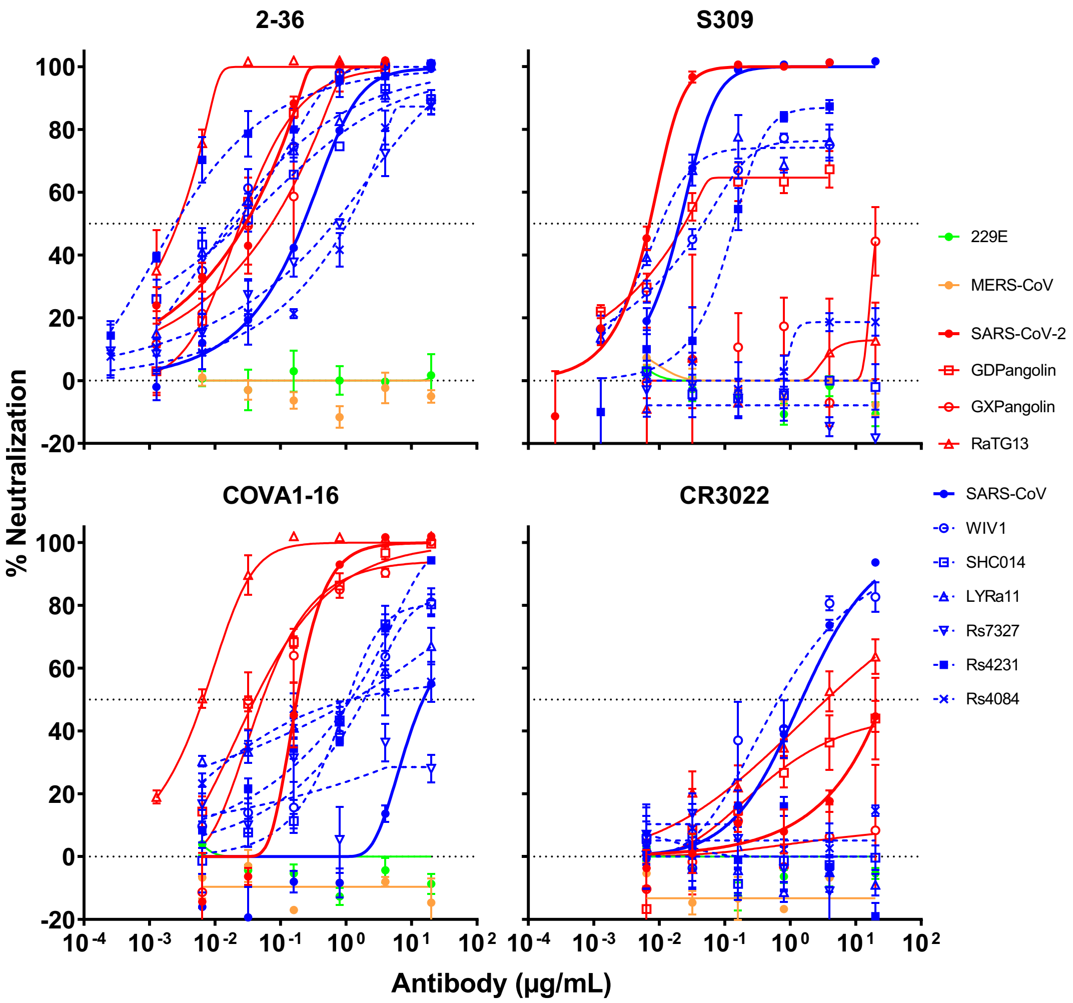


**Supplementary Figure 7.** 2-36 Neutralizes SARS-like coronaviruses using hACE2.

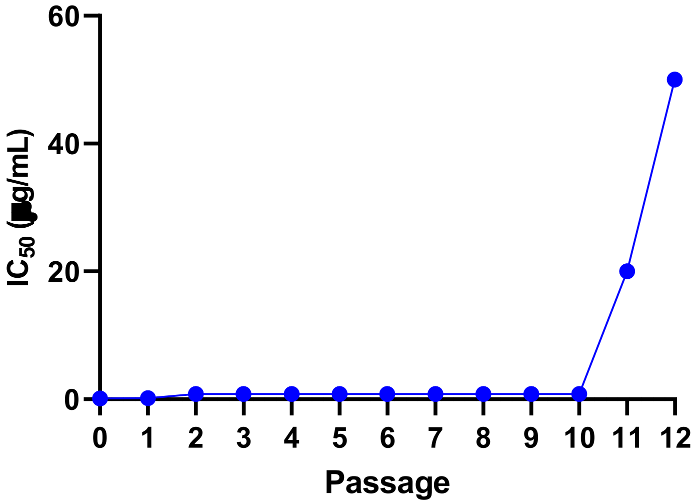


**Supplementary Figure 8.** 2-36 neutralization $IC_{50}$ (µg/mL) on the serially passaged virus.

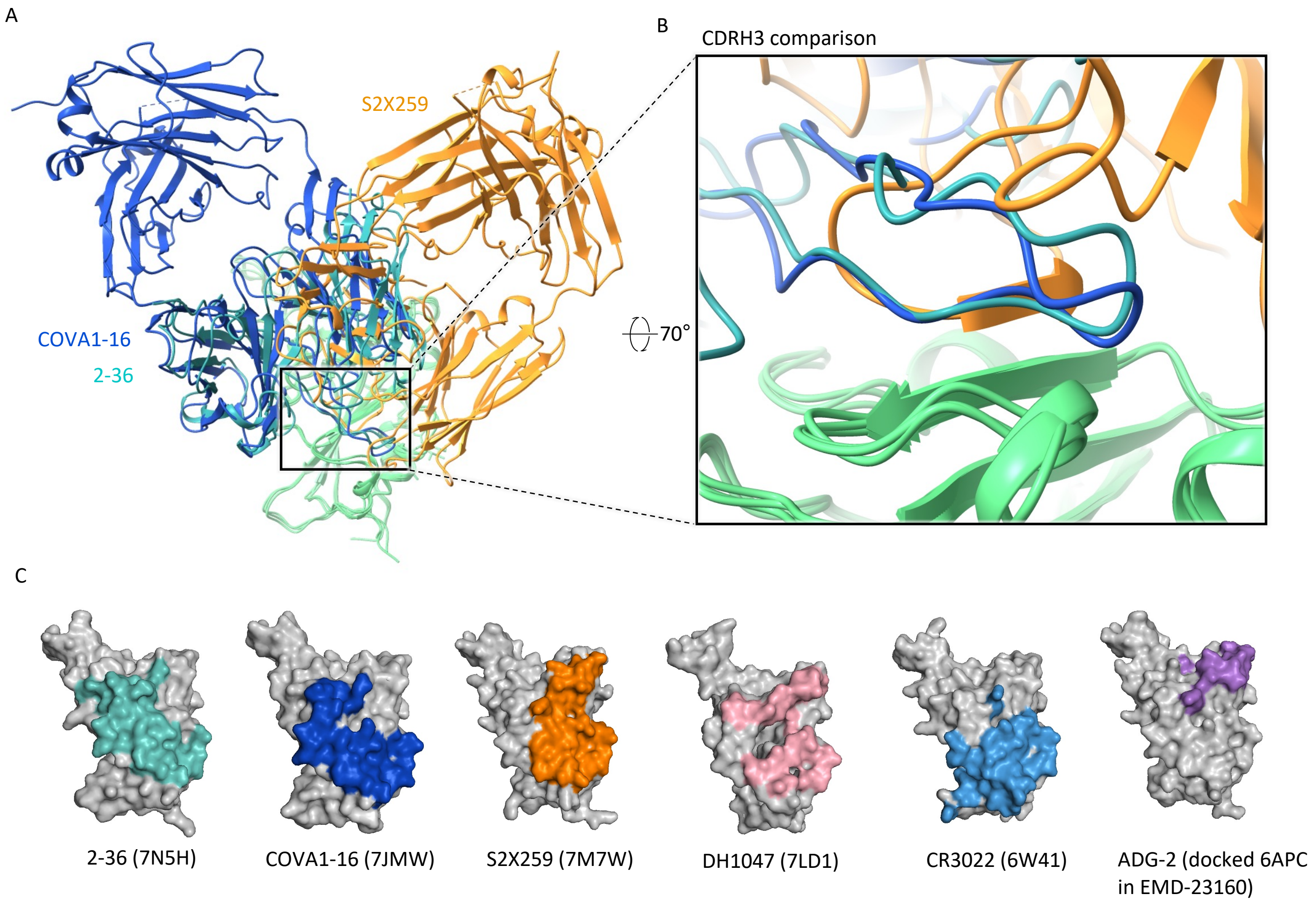


**Supplementary Figure 9. Structural comparison between antibody 2-36 in complex with SARS CoV-2 RBD and other published antibody structures.**

**Supplementary Table 1.** Cryo-EM data collection, processing, and model refinement and validation statistics. Related to Figure 2 and 4.

| | SARS-CoV-2 S2P + 2-36 Fab | SARS-CoV S2P + 2-36 Fab |
|---|---|---|
| **EMDB ID** | 24190 | |
| **PDB ID** | 7N5H | |
| **Data Collection** | | |
| Microscope | FEI Titan Krios | FEI Titan Krios |
| Voltage (keV) | 300 | 300 |
| Magnification | 81000 | 81000 |
| Defocus Range (µm) | -0.8/-2.5 | -0.8/-2.5 |
| Camera | Gatan K3 BioQuantum | Gatan K3 BioQuantum |
| Pixel Size (Å/pix) | 1.07 | 1.07 |
| Recording Mode | counting | counting |
| Dose Rate (e-/pixel/s) | 16 | 16 |
| Elctron Dose (e-/$Å^2$) | 42 | 42 |
| **Data Processing** | | |
| Software | cryoSPARC v2.15 | cryoSPARC v2.15 |
| Micrographs used | 4,226 | 3835 |
| Number of Particles | 171,897 | 80112 |
| Symmetry | C1 | C1 |
| Box Size (pix) | 440 | 440 |
| Global Map $FSC_{0.143}$ (Å) | 3.24 | 4.60 |
| Local Map $FSC_{0.143}$ (Å) | 4.07 | |
| **Refinement and Validation** | | |
| Software | Phenix 1.18 | |
| Initial Model Used | 6BE2 (Fab), 7BZ5 (RBD), 6VXX (spike) | |
| Number of Atoms | 29,691 | |
| Protein Residues | 3,739 | |
| Ligands | NAG: 41 | |
| Model vs. Data CC (mask) | 0.72 | |
| RMS deviations | | |
| Bond lengths (Å) (# >4σ) | 2 | |
| Bond angles (°) (# >4σ) | 17 | |
| MolProbity Score | 1.20 | |
| Clashscore (all atom) | 4.20 | |
| Poor rotamers (%) | 0.18 | |
| Ramachandran plot | | |
| Favored (%) | 98.34 | |
| Allowed (%) | 1.66 | |
| Outliers (%) | 0.00 | |
